# Membrane-Bound O-Acyltransferase 7 (MBOAT7) is a Key Regulator of Glycolysis in Clear Cell Renal Carcinoma

**DOI:** 10.1101/2020.02.10.942458

**Authors:** Chase K. A. Neumann, William Massey, Danny Orabi, Renliang Zhang, Daniel J. Silver, Justin D. Lathia, J. Mark Brown

## Abstract

**Objective:** The most common and deadliest urological cancer is clear cell Renal Cell Carcinoma (ccRCC). ccRCC is characterized by striking reorganization of both carbohydrate and lipid metabolism. It was recently demonstrated that lipid remodeling enzyme Membrane-Bound O-Acyltransferase 7 (MBOAT7) that generates phosphatidylinositol (PI) is important for the ccRCC progression. However, whether MBOAT7-driven PI remodeling is associated with other metabolic alterations commonly found in ccRCC is poorly understood.

**Methods:** MBOAT7 deficient ccRCC cell lines were generated by genome editing, and were characterized by a general reduction in glycolytic capactiy. Using targeted metabolomics approach in Caki-1 cells, we measured the glycolytic intermediates and the relative expression of key glycolytic enzymes. We also measured basal respiration and maximal respiration with MBOAT7 deficiency in the presence of glucose. Lastly, in vivo xenograft studies were performed with parental and MBOAT7 deficient cells.

**Results:** MBOAT7 deficiency was associated with a reduction in glycolytic gene expression and protein abundance. In parallel, we found that glycolytic intermediates similarly decreased which may contribute to a reduction in glycolysis. MBOAT7 deficiency reduces basal respiration and maximal respiration. Similarly, we see maximum glycolytic capacity also reduced with MBOAT7 loss of function. Finally, the in vivo xenograft demonstrated MBOAT7 knockout significantly increased overall survival and reduced glycolytic *HK2* protein abundance *in vivo*.

**Conclusions:** Our work highlights MBOAT7 as a key regulator of glycolysis in ccRCC. Our data provides additional evidence that suggests MBOAT7 as a novel target to regulate tumor growth *in vivo*.

## 1. Introduction

The most prevalent and devastating kidney cancer is clear cell Renal cell carcinoma (ccRCC) [1]. ccRCC has a large lipid rich tumor component compared to other solid tumors [2–4]. Several studies have demonstrated a lipid droplet protein (*PLIN2*) is a biomarker of ccRCC tumors [4,5]. Past studies in other models have shown, signaling phospholipids such as phosphatidylinositol and phosphoinositides (PIPs) are critical for life [6,7]. Previous work has demonstrated that PI3K and PIP_3,4,5_ is critical for Aldolase mobilization to facilitate glycolysis [8]. This work demonstrates a clear link between phosphoinositide signaling and glycolytic regulation.

Similar to many other cancers, ccRCC is characterized by high glycolytic rates that support Warburg metabolism [9]. With dependency on glycolysis, ccRCC upregulates glucose transporter *GLUT1*, increases the rate-limiting hexokinase II (*HK2*), and several other members of glycolysis through hypoxia inducible factors (HIF). Although it is appreciated that glucose metabolism is altered in ccRCC renal physiology upon transformation, the mechanisms of modulating glucose metabolism following transformation is less understood [10].

Furthermore, our recent work demonstrated arachidonic acid-containing phosphatidylinositol (AA-PI) and Lands’ Cycle remodeling enzyme, *MBOAT7*, is important for tumor PI generation, downstream signaling and tumor formation (Neumann et al. 2020. Mol. Metab. *In Press*). However, the mechanism by which MBOAT7 altered tumor biology was not completely understood. *MBOAT7* alters specifically PI and is not promiscuous across other phospholipid species [11,12]. *MBOAT7* knockout has been shown to decrease phosphoinositide levels in multiple tissues through decreasing substrate availability [12,13]. A key ccRCC cell line used in this study is 786-O; this line has a known *PTEN* mutation which confers a more aggressive and metabolically active phenotype *in vivo* and *in vitro [14,15]*. Although lipid metabolism and ccRCC has recently been studied [16], little has been done to address the relationship between lipid metabolism and glycolysis. Here we demonstrate that *MBOAT7* deficiency decreases glycolysis through Aldolase and *HK2*. MBOAT7 loss of function reduces mitochondrial respiration and ATP production, which results in a dramatic increased survival *in vivo* in the 786-O xenograft model. These data support the hypothesis that limiting phosphoinositide phospholipids reduces glycolysis and may hold therapeutic promise.

## 2. Methods

### 2.1 Cell Lines and Cell Culture Conditions

Caki-1 and 786-O cell lines were obtained from American Type Culture Collection (ATCC), and were confirmed to be mycoplasma free. These cells were maintained in McCoy’s 5A Medium with 10% FBS and 1% Pen/Strep. Confirmed *MBOAT7* deficient clones were utilized from previous studies (Neumann et al. 2020. Mol. Metab. In Press). Caki-1 and 786-O expansions were propagated no longer than eighteen consecutive passages.

### 2.2 RNA isolation, quantitative real time-PCR, and RNAseq dataset

Total RNA was isolated the RNeasy isolation method using manufacturer’s recommendations (Qiagen and Thermo Fisher). RNA concentrations were quantified using a Nanodrop 2000. mRNA expression levels were calculated based on the ΔΔ-CT method. Quantitative real time- PCR (qRT-PCR) was conducted using the Applied Biosystems 7500 Real-Time PCR System. Primers used for qRT-PCR are available in Table 1. We used the RNAseq dataset previously published from the Caki-1 cell line available with the GEO accession: GSE131881 (Neumann et al. 2020. Mol. Metab. In Press).

**Table 1.**
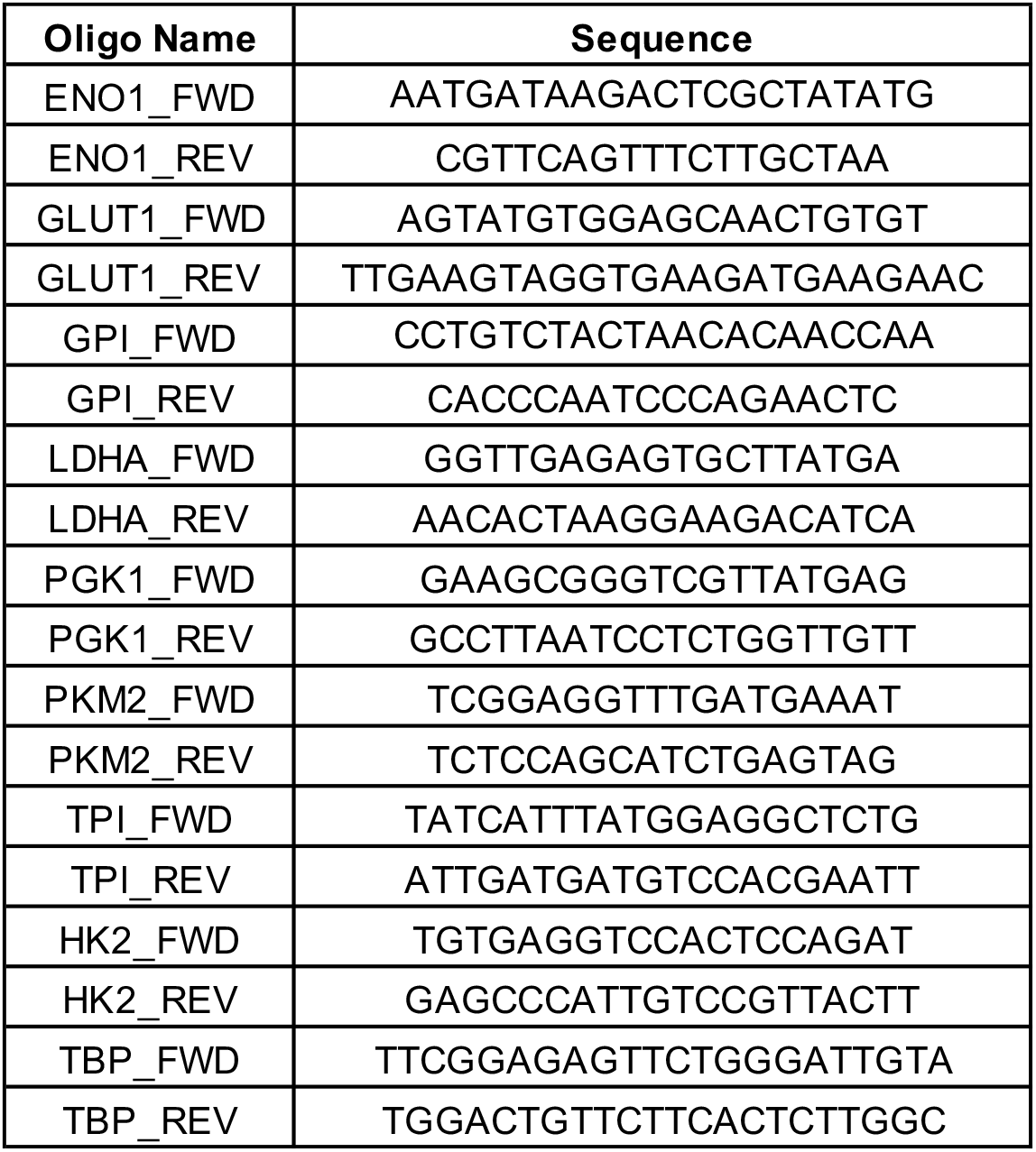
Quantitative Real Time – PCR Primer Sequences

### 2.3 Western Blot Analysis

Cell tissue lysates were generated using a modified RIPA buffer and western blotting was performed following the previously described methods [17]. Antibodies used are available through Sigma Aldrich or Cell Signaling with the corresponding product numbers in Table 2.

**Table 2.**
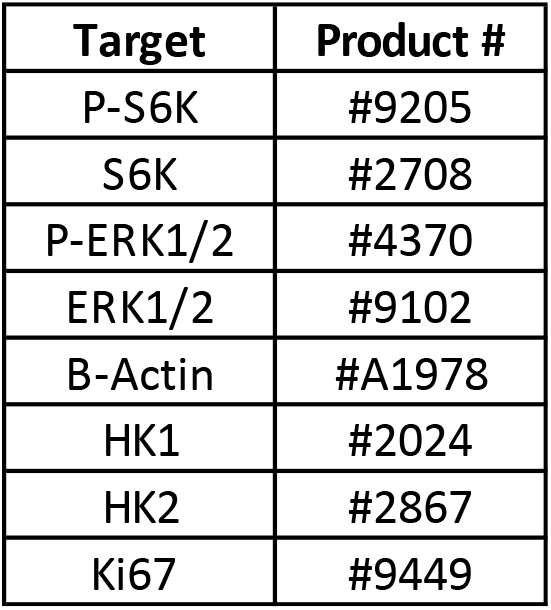
Western Blotting and Immunohistochemistry Antibodies

### 2.4 Seahorse Metabolic Fitness Studies

Following the Agilent Seahorse manufacturers recommended protocol, we plated cell populations into XFe24 culture plates the day before assay at 50,000 cells per well in normal growth media (McCoy’s 5A). Both Caki-1 and 786-O were used at the same cell density. *Mitochondrial Stress Test:* The media was changed from growth media to assay media 1 hour prior to testing. DMEM Agilent base media + 10mM glucose for the mitochondrial stress test was used for the initial assays. The traditional Mito Stress Test assay was used with all studies Oligomycin 1uM (Port A), FCCP 1uM (Port B), and Rot/AA 0.5 uM (Port C). *Glycolysis Stress Test:* The media was changed from growth media to base media or base media + 1mM pyruvate at 1 hour prior to testing. Following manufacturers recommendations Glucose 10mM (Port A), Oligomycin 1uM (Port B), and 2-DG 50mM (Port C). Both Glycolysis and Mitochondrial Stress Test were normalized to well protein concentrations to account for variability in cell number.

### 2.5 In Vivo ccRCC Xenograft Studies

MBOAT7^+/+^ and MBOAT7^−/−^ 786-O cell lines were injected in the subcutaneous flank of NSG animals (Jackson laboratory) at 2.5 million cells per mouse in PBS (n=10 per group). Once tumors were palpable, digital caliper measurements were used to follow tumor growth over time. Endpoint criteria were defined as 15 mm tumor width or length, these animals were then necropsied and reached endpoint. All mice studies were approved by the Institutional Animal Care and Use Committee of the Cleveland Clinic.

### 2.6 Statistical Analysis

All figures are shown with ± SEM. For comparisons of three groups, we utilized one-way ANOVA with a *post hoc* Tukey test. Log-rank test (Mantel- Cox test) was used to compare survival differences between two groups of patients. When comparing two groups, we used an unpaired Student t test or multiple t test. The P values significance cutoffs for all tests used are as follows: p-value < 0.05(*), 0.002(**), 0.0002(***), <0.0001(****).

## 3. Results

### Loss of MBOAT7 correlates with a decrease in glucose consumption and glycolytic gene expression

To study the role of MBOAT7 and PI metabolism on glycolysis, we utilized previously published MBOAT7 deficient ccRCC Caki-1 and 786-O cell. We observed that with MBOAT7 loss of function, the acidification of media occurs more slowly compared to wild-type (Figure 1A). Utilizing a previously described RNAseq dataset in the Caki-1 cell line (GSE131881), we see that MBOAT7 deficiency leads to a significant reduction in the glycolytic gene set after gene set enrichment analysis (GSEA) (Figure 1B). Looking specifically at the initial steps of glycolysis, there is a reduction in the hexokinase 2 (*HK2*), glucose-6-phosphate isomerase (*GPI*), and phosphofructokinase (*PFKP*) with MBOAT7 loss of function in the RNAseq dataset (Figure 1C). *In vitro* experiments, we see that MBOAT7 deficiency leads to a decrease in glucose consumption and lactate production (Figure 1D), consistent with decreased glycolysis.

**Figure 1.**
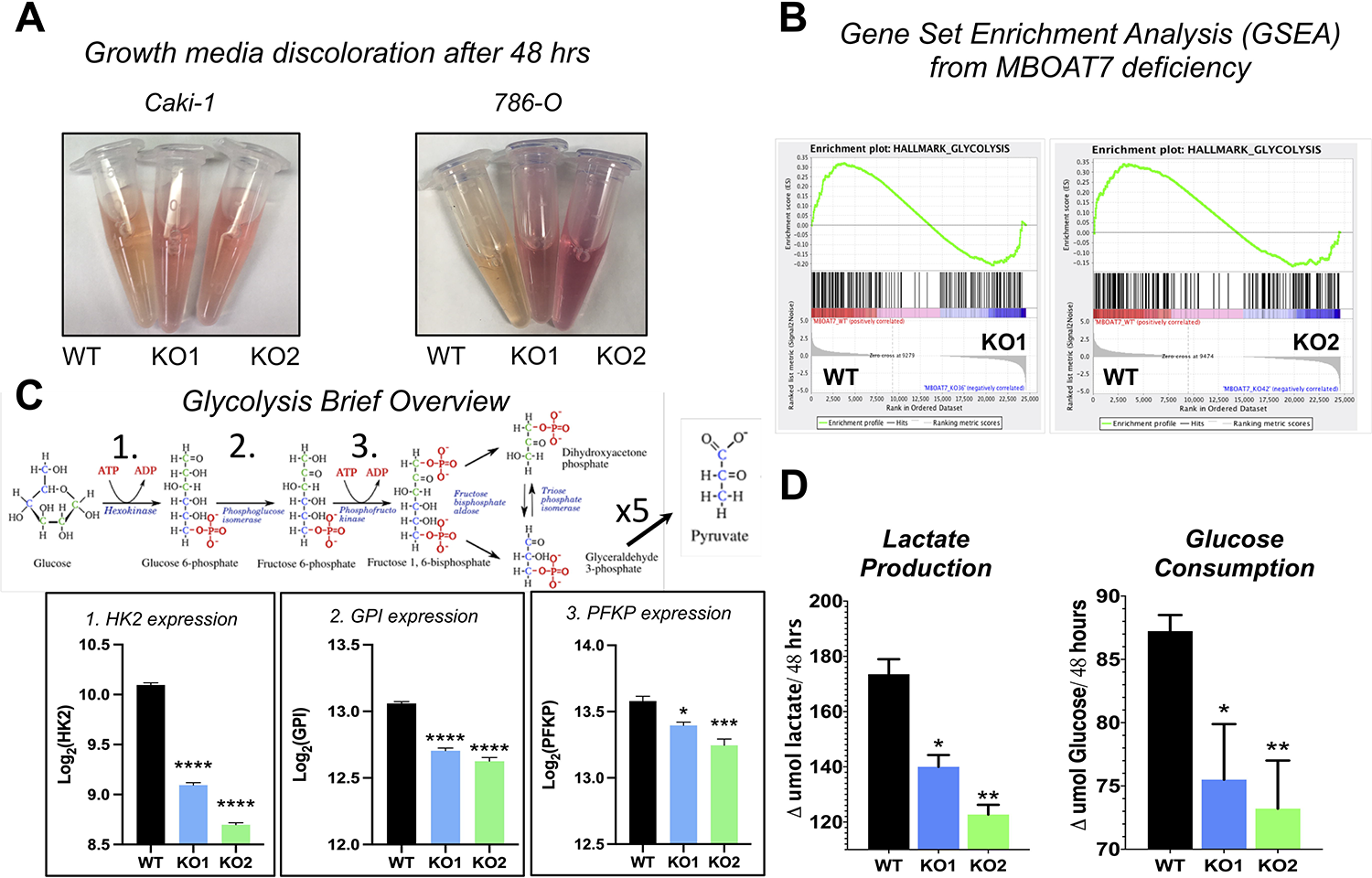
MBOAT7 deficiency correlates with a reduction in glycolysis. **A)** Conditioned media from cells seeded at the same original density and after 48hrs of culture. **B)** Gene Set Enrichment Analysis (GSEA) of RNAseq dataset demonstrates a reduction in glycolysis enrichment with MBOAT7 loss of function. **C)** The first three steps of glycolysis demonstrate a significant reduction with MBOAT7 deficiency. **D)** Using the supernatant of from the MBOAT7 WT and KO cell lines following 48hrs of media, glucose consumption measurements decrease with loss of function (n=4 per group). Similarly, lactate production measurements after 48hrs decrease with MBOAT7 deficiency (n=4 per group). Student t-test: * < 0.05, ** < 0.0021, *** < 0.0002

### MBOAT7 deficiency reduces pyruvate and several key glycolytic enzymes in HK2 and ALDOC

We pursued targeted mass spectrometry for the glycolytic intermediates in the Caki-1 model following high glucose supplementation for 1 hour. Dihydroxyacetone and pyruvate, an intermediate and product of glycolysis (Figure 2A) demonstrates a significant reduction following *MBOAT7* loss of function, which corresponds with the loss of gene expression in *HK2* and *ALDOC (Figure 2B)*. To validate the reduction in *HK2* expression, we performed western blotting and see that HK2 protein abundance decreases with MBOAT7 deficiency in the Caki-1 and 786-O models (Figure 2C). These data suggest alterations in glycolysis and metabolic fitness with MBOAT7 deficiency.

**Figure 2.**
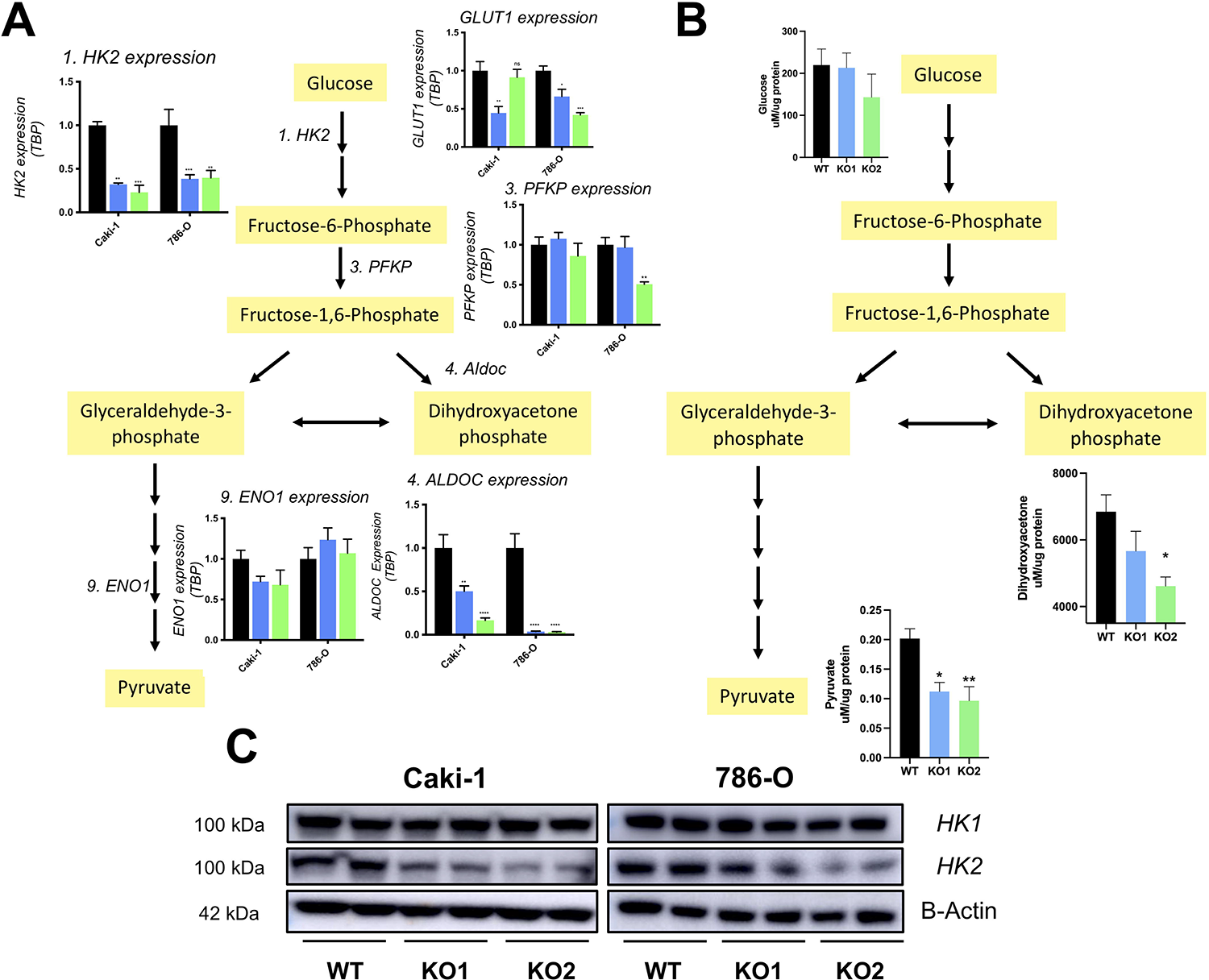
MBOAT7 deficiency decreases glycolytic product, pyruvate. **A)** Quantitative Real Time PCR (qRT-PCR) shows reduced glycolytic gene expression specifically, 1. *HK2* and 4. *ALDOC.* (n=5 per group) **B)** Targeted mass spectrometry of three glycolytic intermediates reveals a reduction in pyruvate with MBOAT7 deficiency. (n=4 per group) **C)** Western blot analysis of *HK1*, *HK2, and B-Actin* in ccRCC cell lines, Caki-1 and 786-O with or without MBOAT7 deficiency. Student t-test: * < 0.05, ** < 0.0021, *** < 0.0002

### MBOAT7 loss of function reduces glycolysis and metabolic fitness

To assess glucose metabolism, we utilized the Seahorse Metabolic Stress Tests. Oxygen consumption rate (OCR) is directly related to ATP production through the electron transport chain. In our initial tests with glucose, MBOAT7 deficient Caki-1 cells have a reduced basal and maximal respiration (Figure 3A). Our findings are validated in the 786-O RCC model that basal and maximal respiration are reduced with the loss of *MBOAT7* (Figure 3B). MBOAT7 deficiency significantly reduces ATP production in the Caki-1 and 786-O cell models (Figure 3C,D). To assess glucose utilization, we performed glycolysis stress test in the presence of base media alone, which demonstrated a reduction in maximum glycolysis with MBOAT7 deficiency (Figure 3E). This significant reduction in glycolysis could be rescued with pyruvate supplementation into the media (Figure 3F).

**Figure 3:**
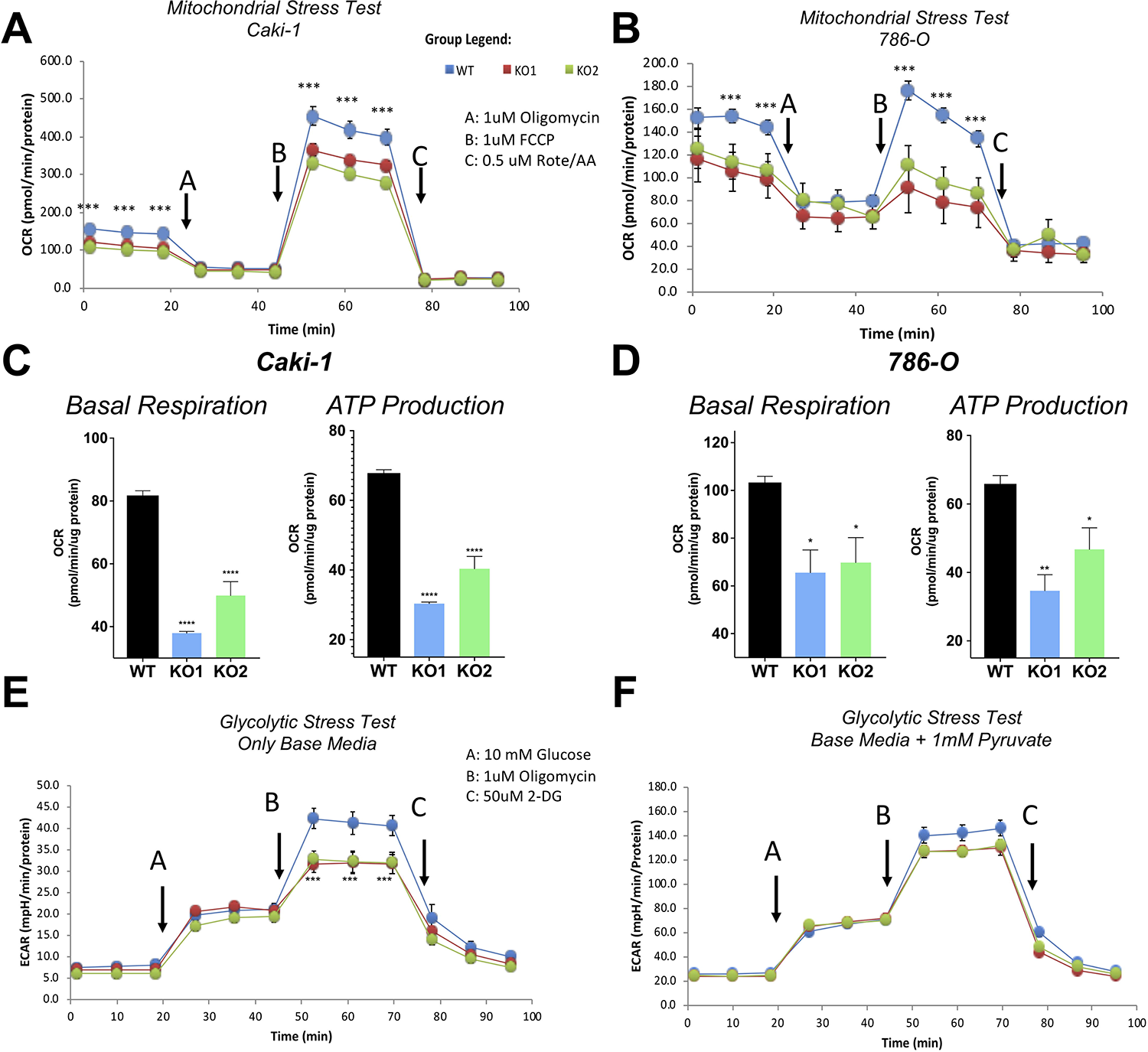
Metabolic fitness and glucose utilization decreases with the loss of MBOAT7. **A)** Mitochondrial stress test with Caki-1 WT, KO1, and KO2 in the presence of glucose. **B)** Mitochondrial stress test with 786-O WT, KO1, and KO2 in the presence of glucose. **C)** Basal respiration and ATP production during mitochondrial stress test in Caki- 1 WT, KO1, and KO2. **D)** Basal respiration and ATP production during mitochondrial stress test in 786-O WT, KO1, and KO2. **E)** Glycolytic Stress Test in the presence of only glucose with Caki-1 cell lines. **F)** Glycolytic Stress Test in the presence of pyruvate and glucose with Caki-1 cell lines rescues maximal glycolytic capacity. (n=5 per group) (2 = independent replicate experiments) Student t-test: * < 0.05, ** < 0.0021, *** < 0.0002

### MBOAT7 loss of function decreases in vivo tumor volume and glycolysis

To test the overall tumor growth, MBOAT7 wild type or deficient 786-O cells were injected into the flanks of Nod SCID Gamma (NSG) mice. In this experiment, we see an increased *in vivo* overall survival with MBOAT7 loss of function (Figure 4A). Accordingly, the tumor volume at end-point for all animals is substantially reduced with MBOAT7 deficiency (Figure 4B). To confirm these data, we utilized histology and immunohistochemistry to stain for the H&E (architecture), Ki67 (cell division), and Hexokinase 2 (glycolysis) (Figure 4C). The quantification of tumor staining with or without MBOAT7 demonstrates a reduction in Ki67^+^ staining per field and trending reductions in *HK2* staining (Figure 4D). The *in vivo* gene expression shows a significant reduction in the rate-limiting glycolytic pathway member, *HK2*, within MBOAT7 deficient tumors (Figure 4E). Other metabolic regulators were assessed in these tumors such as CPT1A, PPARA, and PPARG **(Supplementary).** Additionally, we also see trending increases in Trichrome fibrosis staining in the MBOAT7 wild-type tumors **(Supplementary).** Mechanistically, MBOAT7 deficiency leads to a decrease in *in vivo* P-S6K and P-ERK signaling, while maintaining reduced *HK2* protein abundance (Figure 4F).

**Figure 4:**
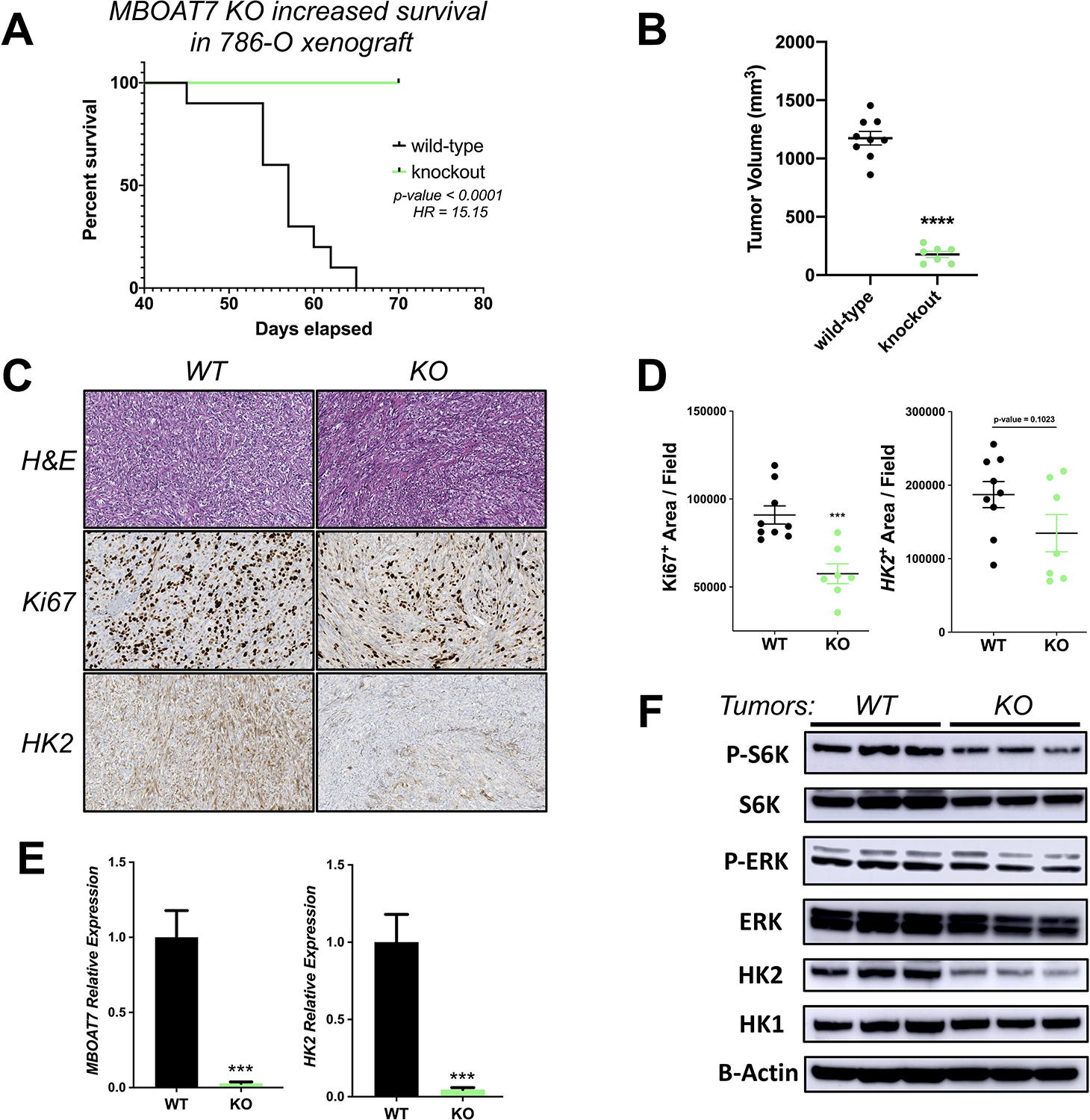
Decreased metabolic fitness and MBOAT7 deficiency leads to increased survival *in vivo*. **A)** 786-O xenografts in NSG female mice show a striking survival advantage with MBOAT7 deficiency. **B)** Tumor volume of 786-O xenografts after meeting experimental end-point criteria or 70 days post injection. **C)** Histology representative images of 786-O WT and KO including H&E, Ki67, and *HK2.* **D)** Quantification of Ki67^+^ staining and *HK2*^+^ *staining* per field in 786-O xenografts. (n=9 or 7 per group) **E)** Tumor mRNA expression of *MBOAT7* and *HK2. (n=5 per group) **F)*** Western blotting analysis of tumor signaling (S6 Kinase and ERK1/2 signaling) and key glycolysis markers (*HK1* and *HK2*). (n=3 per group)

## 4. Discussion

A hallmark of many cancers including kidney cancer is the unrelenting metabolic adaptation and glycolysis that occurs. To meet metabolic demand, tumor metabolism increases glycolysis, which provides a relatively inefficient ATP production. The Warburg effect in full force leads to increased glycolysis and substrate influx to help fuel the increased anabolism for cell division and proliferation. Glycolytic gene expression can be altered through multiple mechanisms including HIF and mTORC [18–21], the continued signaling through these nodes can lead to increases in glycolytic enzymatic intermediates. Others have shown PI3K and subsequent product (PIP_3,4,5_) is critical for the downstream signaling and increased glycolytic activity through mTORC1/S6K [8,15].

In our previous study that identified MBOAT7’s potential role in cancer, we demonstrated a loss of PI confers a decreased S6K signaling (Neumann et al. 2020. Mol. Metab. In Press). To follow, we see a reduction in glycolysis with MBOAT7 loss of function in both ccRCC cell lines. In this study, we demonstrate in ccRCC that with the loss of a Lands’ Cycle remodeling enzyme, MBOAT7, and the reduction in PI leads to reduced glycolysis. The reduced glycolytic gene expression with MBOAT7 deficiency was validated using qRT-PCR, which showed the most striking reductions in glycolytic gene expression presented in the Hexokinase-2 (*HK2*) and Aldolase (*ALDOC*) transcripts (Figure 2). Pairing gene expression with the glycolytic intermediate profiles, suggests a reduction in glycolysis is occurring as the product of glycolysis, pyruvate, is reduced by approximately 50%.

Consistent with the reduction in glycolysis, the metabolic fitness with MBOAT7 loss of function suggests a reduction in basal respiration and maximal respiration in the presence of only glucose. MBOAT7 deficient ccRCC lines seem to be not utilizing glucose as efficiently compared to wild-type cells given the same constraints. The utilization of glucose for tumor growth and development is incredibly important, and led us to our final experiment.

Previous MBOAT7 loss of function studies utilized the Caki-1 ccRCC cell xenograft, and MBOAT7 deficient xenografts never developed palpable tumors after 9 weeks. In this study, our *in vivo* experiments utilized a more aggressive 786-O model with a *PTEN* mutation, and as the mechanism for *PTEN enriches* in PIP_3_ species. These MBOAT7 deficient xenografts still developed palpable tumors as we hypothesized due to a more aggressive genomic background, however these tumors are delayed with a slower tumor growth and never reach endpoint criteria. At endpoint MBOAT7 deficient tumors were ~10% tumor volume of wildtype 786-O xenografts.

Interestingly, previous studies have shown that PTEN mutations increase basal glycolysis and cell growth of mouse embryonic fibroblast (MEF) [22]. For the first time, this study demonstrates that limiting upstream PI production through MBOAT7 can limit glycolysis, metabolic fitness, and *in vivo* tumor growth regardless of *PTEN* status. This study gives more evidence that PI metabolism helps regulate glycolysis. This also provides rationale for several future studies including assessing MBOAT7/AA-PI role in other PI3K/mTORC1 dependent cancers, *in vivo* MBOAT7 knockdown interventional study with *PTEN*^*mut*^/PI3K/mTORC1 active cancer models, developing small molecule inhibitors towards MBOAT7, and determining the role for Lands’ Cycle in cancer progression.

## Author Contributions

C.K.A.N., J.D.L., J.M.B. planned the project, designed experiments, and wrote the manuscript; C.K.A.N., R.Z., D.J.S., W.M., and D.O. executed experiments. C.K.A.N., R.Z., D.J.S., W.M. and D.O. analyzed the data; J.D.L., and J.M.B. provided financial support; all authors were involved in the editing of the final manuscript.

## Acknowledgments

This study was supported in part by grants provided by the National Institutes of Health (NIH): R01 DK120679 (J.M.B.), P50 AA024333 (J.M.B.), and P01 HL147823 (J.M.B.). Development of glycolytic mass spectrometry methods reported here were supported by generous pilot grants from the Clinical and Translational Science Collaborative of Cleveland (4UL1TR000439) from the National Center for Advancing Translational Sciences component of NIH and the NIH Roadmap for Medical Research, the Case Comprehensive Cancer Center (P30 CA043703), the VeloSano Foundation, and a Cleveland Clinic Research Center of Excellence Award.

